# PathoResist AI: A One-Click Web Platform for Rapid Pathogen Resistance Analysis Based on the all_ratio Algorithm

**DOI:** 10.64898/2026.02.12.705264

**Authors:** Guoqin Mai, Ying Dai

## Abstract

This study introduces a one-stop analysis platform named “PathoResistAI” (https://www.resistpath.com/), which can be used to solve the technical bottlenecks of pathogenic microorganism detection and antimicrobial resistance analysis. The platform is based on nanopore sequencing and the innovative all-ratio algorithm, which integrates four-dimensional parameters (sequence similarity, abundance, matching number, and matching length), significantly improving the detection accuracy of low-abundance pathogens and drug-resistance genes. The platform adopts four layers of modular design (input layer, core engine, dual-channel output, and visualization layer). Users only need to upload data in FASTQ format, and they can obtain automated reports, including pathogen identification and drug-resistance gene prediction within 30 min. The verification results show that the platform can accurately identify bacteria (e.g., *Staphylococcus aureus* and *Serratia marcescens*), viruses (e.g., Ebola virus), and drug-resistance genes (e.g., SdeY), which are consistent with the published literature results. Limitations include only supporting long-read sequencing data, small sample size (fewer than 50 cases), and lack of real clinical sample verification. In general, this platform represents the application and exploration of nanopore sequencing in the field of rapid detection of pathogenic microorganisms, and provides a new tool for microbial pathogen or AMR detection research.

## Introduction

Infectious diseases caused by infectious pathogens pose a major threat to global health and cause serious economic burden [1]. Antimicrobial resistance (AMR) has become a global public health crisis. According to the World Health Organization report from 2024, AMR directly causes at least 1.27 million deaths every year, and it is expected to cause trillions of dollars in economic losses by 2050 [2]. The existing detection technology faces three core bottlenecks. First, the timeliness is poor. For traditional PCR, pathogens have to be cultivated and isolated, which lead to the lag in clinical decision-making [3]. Second, the detection dimension is single, and simultaneous pathogen typing and drug-resistance gene spectrum identification is rarely possible [4]. Third, the sensitivity is insufficient, and the rate of missing detection of low-abundance drug-resistant bacteria in complex samples such as blood is high [5].

This technological gap has spawned the demand for a new generation of integrated detection solutions. It is necessary to complete the whole process analysis from sample to report within 6 h and realize pathogen identification (such as carbapenemase-*Enterobacter aerogenes*) and drug-resistance mechanism analysis (such as blaKPC gene detection). Nanopore sequencing technology has real-time advantages, and it has been applied for the detection of various pathogenic microorganisms, such as bacteria, fungi, viruses, and parasites [6-10]. However, nanopore sequencing technology has high requirements for samples [5], and its original data have a certain error rate [11], which seriously restricts its clinical applicability. The newly researched all_ratio algorithm [12] provides a multidimensional comprehensive evaluation framework by integrating four key parameters—sequence similarity, abundance, matching number, and matching length—which can more accurately reflect the identification reliability of pathogenic microorganisms and AMR than a single parameter. Specifically, in complex samples, a single parameter may have the risk of misjudgment, whereas the all_ratio algorithm effectively reduces the false-positive and false-negative rates by weighted fusion of multiple parameters, improving the accuracy of pathogen and AMR identification.

These advantages make the all_ratio algorithm a potential auxiliary method for the detection of pathogenic microorganisms or AMR. However, to apply the all_ratio algorithm, it is necessary to download relevant databases and to download and install relevant software. This is a little cumbersome so this algorithm is not conducive for use by novice researchers. Therefore, in this study, we developed a one-click submission platform based on the all_ratio algorithm, which is easy to operate and learn. Users only need to submit nanopore (long-reads) sequencing data on the website. About 30 min later, the table and figure results of suspected pathogenic microorganisms or antibiotic-resistance genes are sent to the user’s email.

## Methods

### 1. Algorithm innovation: The all_ratio quadruple-weight design

The core breakthrough of the all_ratio algorithm [12] lies in the dynamic fusion of four types of bioinformatics parameters, namely sequence similarity, abundance, matching number, and matching length, and the synchronous and accurate detection of pathogens and drug-resistance genes.

### 2. Platform architecture: Four-layer modular design

The platform is based on the all_ratio algorithm of Maiguoqin et al. [12]. Maiguoqin designed the website and provided the all_ratio algorithm (https://github.com/guoqinmai/nanopore_code_data_4_v2/tree/main/code). The prediction platform PathoResistAI was developed by the third-party technical team Fujian susuzhao Technology Co., Ltd. (Xianfeng building, No. 1 Futian Road, Fengze District, Quanzhou, Fujian, China), based on the all_ratio algorithm. The platform mainly includes four layers of modular design, namely an input layer, the core engine, a dual-channel output, and a visualization layer (Figure 1).

**Figure 1.**
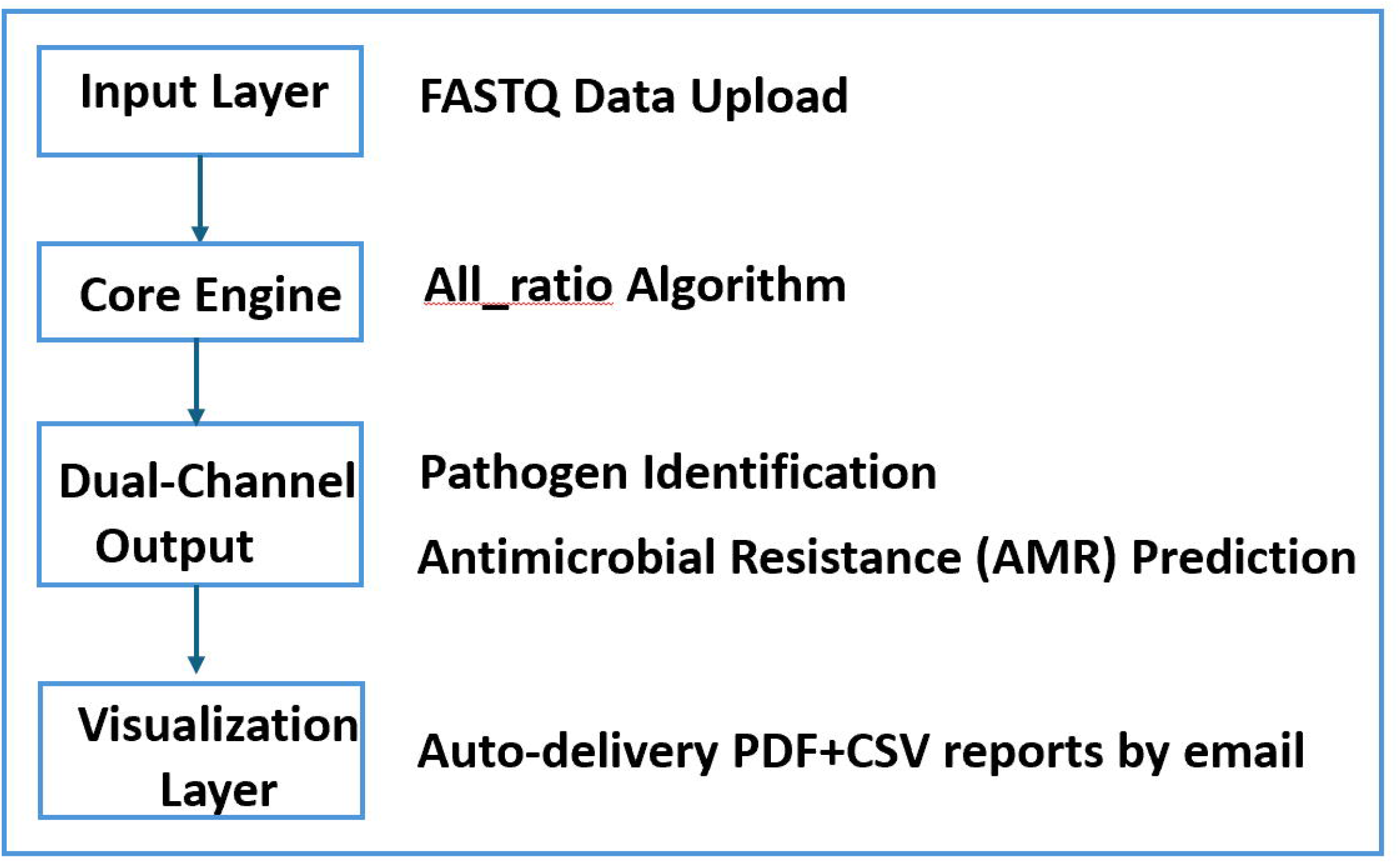
Technology roadmap of the platform “PathoResist AI.” The platform mainly includes four layers of modular design, namely an input layer, the core engine, a dual-channel output, and a visualization layer.

#### (1) Input layer

The input data support the FASTQ format of nanopore (long-reads) sequencing data, which is easy to learn and can be submitted with one click. There is a progress bar to observe the upload status. Moreover, the user needs to fill in the sample number and email address to prepare for receiving the results. An example of input is available for download on this page (Figures 1 and 2).

**Figure 2.**
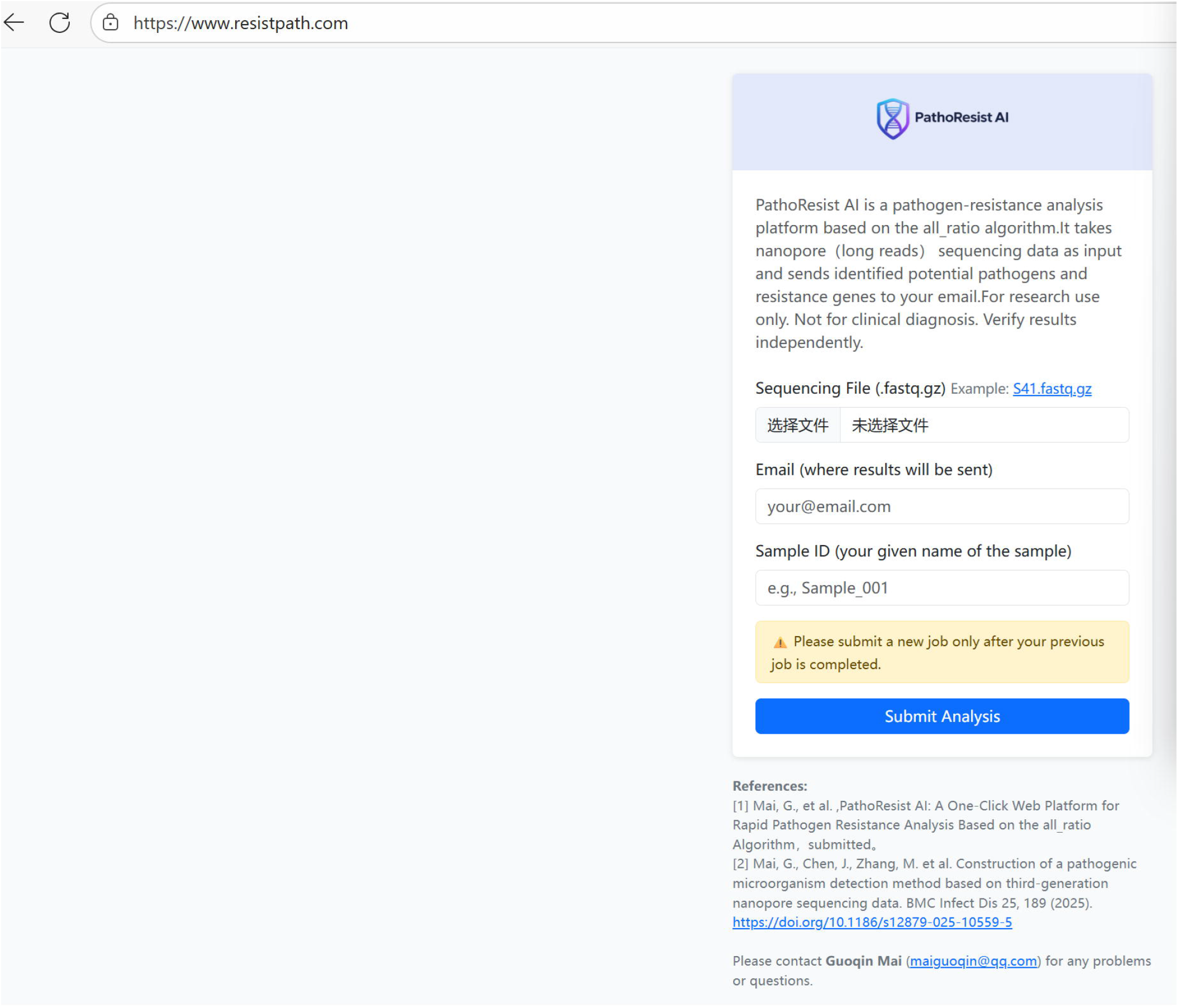
Interface of the platform “PathoResist AI.” The PathoResist AI is a pathogen–resistance analysis platform based on the all_ratio algorithm. It takes nanopore (long-reads) sequencing data as input and sends identified potential pathogens and resistance genes to the provided email address. It is suitable for research use only and not for clinical diagnosis. The results should be verified independently.

#### (2) Core engine

The all_ratio algorithm calculation flow, including quality control, removal of host pollution, and BLASTN, calculates the comprehensive parameters of four-dimensional parameter fusion, such as all_ratio [12], which are automatically run in the background (Figure 1).

#### (3) Dual-channel output

Pathogen identification, based on the database of bacteria, fungi, viruses, and parasites, includes 8565 microbial nucleic acids [12]. Drug-resistance prediction is based on AMRFinderPlus/database, which includes 5956 antibiotic-resistance genes [13] (Figure 1).

#### (4) Visualization layer

The table with detailed information of pathogen identification or antibiotic-resistance gene identification includes six parameters, namely number_of_matches, average, sum, reads_ratio, all_sum, and all_ratio [12]. If bacteria are identified, in addition to the information table of pathogen identification, the table of drug-resistance gene identification and its visual box plot are generated. After uploading the data successfully, the relevant results are automatically sent to the user’s email address within about 30 min (Figures 1 and 2).

### 3. Validation data

In this study, we used the nanopore sequencing data of clinical samples from published articles [14, 15] to verify the performance, which can detect pathogens and antibiotic-resistance genes at the same time, and compared the obtained results with the results reported in the published articles. The timeliness test was also conducted. The average time from FASTQ to the full report was 30 min (background server configuration: CPU 128, MEM 1024G). Generally, the larger the amount of uploaded data, the longer the running time required.

## Results

The detection results of sample S6 (Figure 3, Supplementary Table 1) showed that the maximum value of the parameter all_ratio of the pathogenic microorganism was 0.97, with its reads_ratio being 0.16, corresponding to the pathogenic microorganism *Staphylococcus aureus*, which is consistent with the published results [14]. The first two maximum values of the parameter all_ratio of antibiotic-resistance genes in sample S6 were 0.53 and 0.31 (Figure 3, Supplementary Table 2), with the reads_ratio being 0.17 and 0.26, corresponding to the antibiotic-resistance genes BlaR1 and CadD, respectively. Literature review revealed that these genes are common in drug-resistant *S. aureus*[16, 17]. Sample S6 had a data file of about 36.67 MB. After uploading the data successfully, it took about 7 min to receive the results by email.

**Figure 3.**
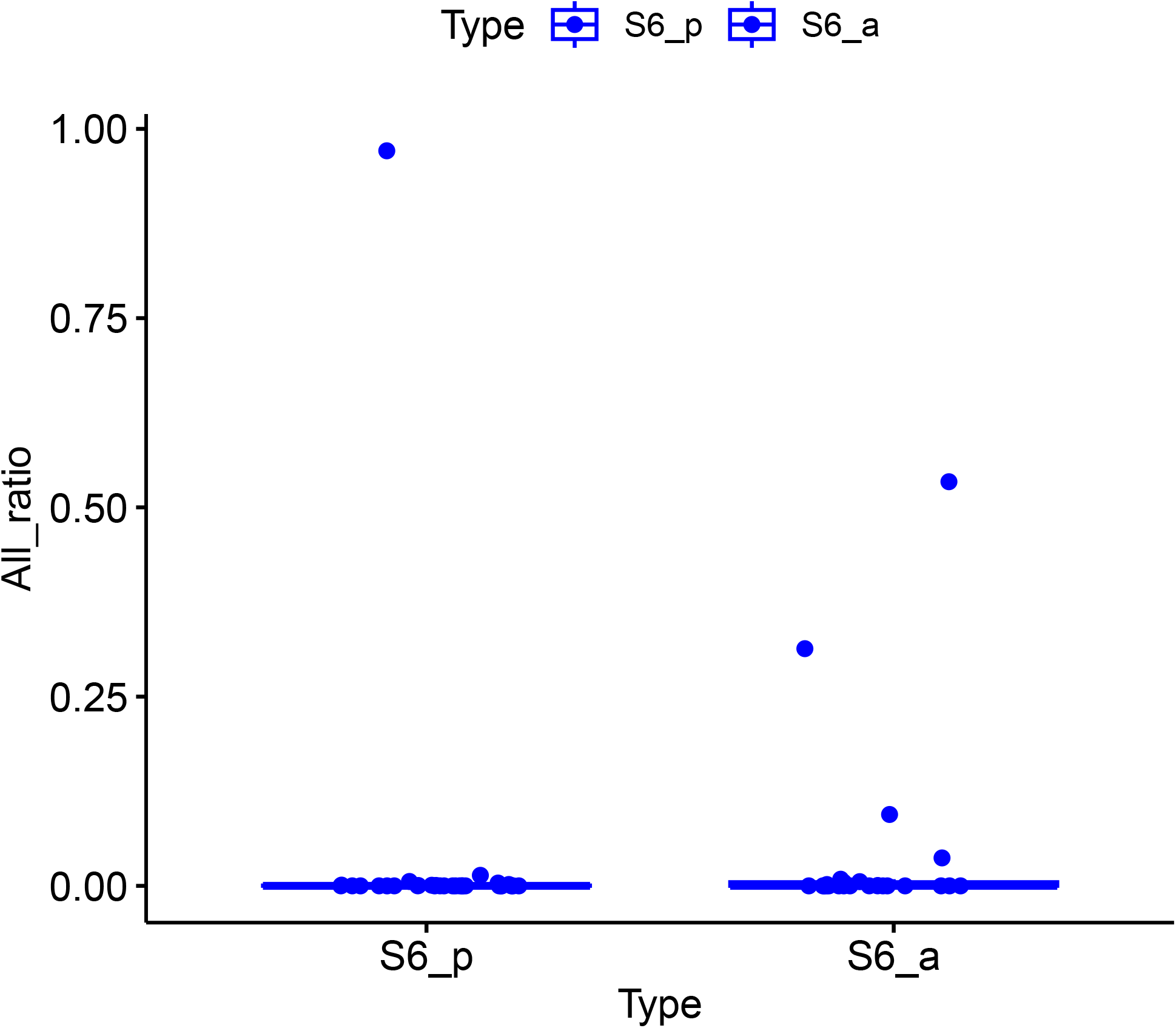
Detection results of sample S6 with detected bacteria and antibiotic-resistance genes.

The detection results of sample S13 (Figure 4, Supplementary Table 3) showed that the maximum value of the parameter all_ratio of the pathogenic microorganism was 0.97, with the reads_ratio being 0.24, corresponding to the pathogenic microorganism *Serratia marcescens*, which is consistent with the published results [14]. The maximum value of the parameter all_ratio of the antibiotic-resistance gene of sample S13 was 0.97 (Figure 4, Supplementary Table 4), with the reads_ratio being 0.20, corresponding to the antibiotic-resistance gene sdeY, which is consistent with the published results [14]. These findings demonstrate that the all_ratio from the website can detect bacteria and antibiotic-resistance genes at the same time. For sample S13, the data file was about 77.67 MB. After uploading the data successfully, it took about 8 min to receive the results by email.

**Figure 4.**
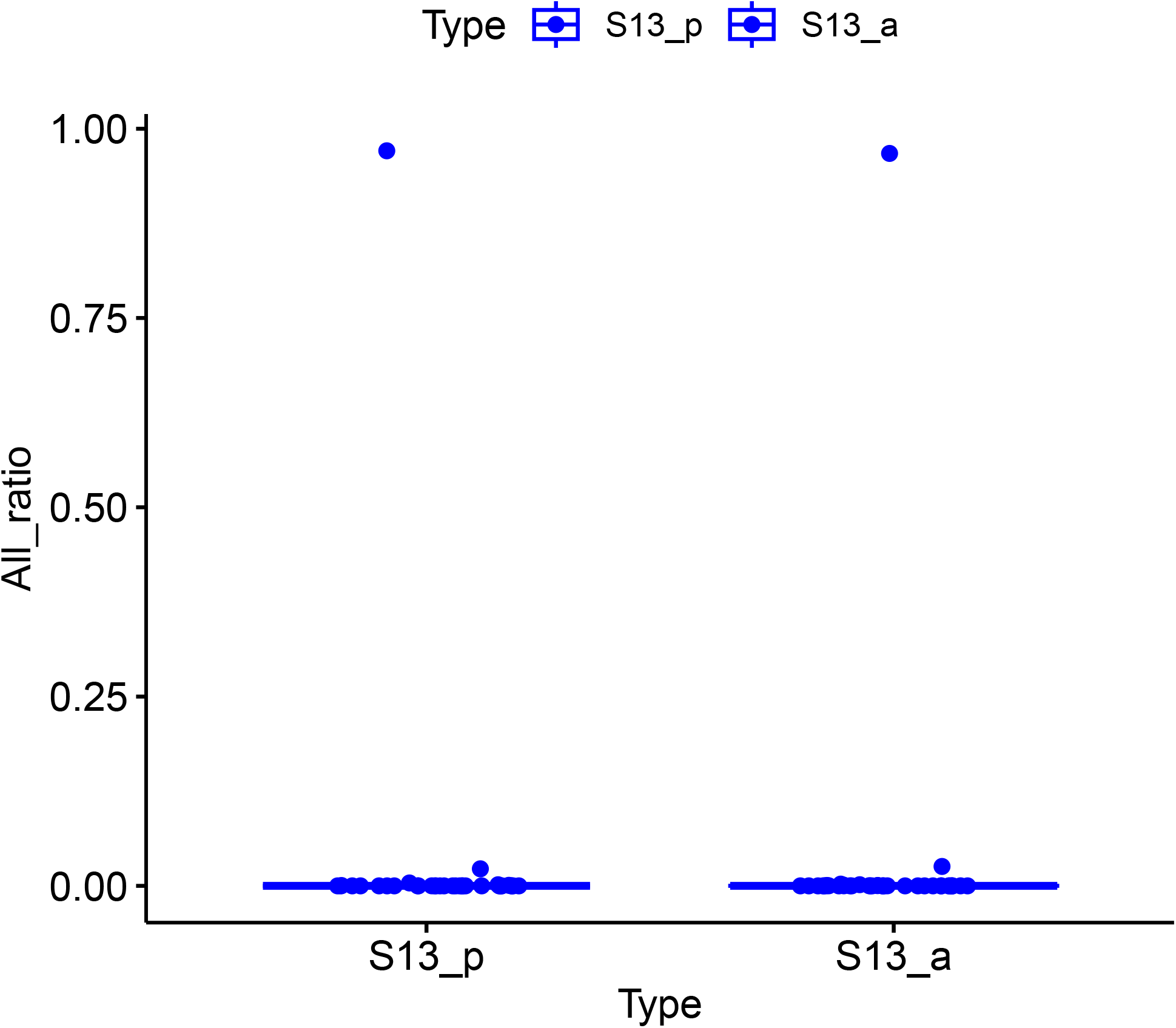
Detection results of sample S13 with detected bacteria and antibiotic-resistance genes.

The platform can also detect virus samples. The maximum value of the parameter all_ratio of sample E1 was 1 (Supplementary Table 5), with the reads_ratio being 1, corresponding to the pathogenic microorganism chikungunya virus, which is consistent with the published results [15]. For sample E1, the data file was about 2.84 MB. After uploading the data successfully, it took about 4 min to receive the results by email. The maximum value of the parameter all_ratio of sample E2 was 0.99 (Supplementary Table 6), with the reads_ratio being 0.30, which corresponds to the pathogenic microorganism Ebola virus, which is consistent with the published results [15], indicating that all_ratio can also detect viruses. For sample E2, the data file was about 16.23 MB. After uploading the data successfully, it took about 4 min to receive the results by email.

## Discussion

The PathoResist AI platform greatly reduces the threshold of grass-roots research through “one-click analysis.” Traditional drug-resistance analysis requires bioinformatics professionals to operate a variety of software tools (such as Kraken2 [18] and ARIBA [19]), while this platform integrates the whole process from original data cleaning and pathogen identification to drug-resistance gene prediction. Users only need to upload nanopore sequencing data to obtain structured reports within 30 min (Figure 1), without coding experience.

The core innovation of the platform is the collaborative design of the all_ratio algorithm and dynamic threshold. Compared with the mainstream tool AMRPlusPlus [20] (depending on the static threshold), the all_ratio algorithm significantly improves the detection rate of low-abundance pathogens by multiplying and weighting the similarity, abundance, matching number, and matching length in real time [12]. The application potential is reflected in the mobile detection scenario. Combined with the portability of MinION devices, the platform can be extended to clinical medicine, public health, animal husbandry and environmental drug-resistance gene screening in the future.

However, our research has some limitations. First, the input data were only long-read sequencing data, such as nanopore sequencing data, and not short-read sequencing data. Second, the sample size was small, given that the all_ratio algorithm and its website tested fewer than 50 samples from the published literature [12]. Therefore, more samples need to be collected in the future to verify the effectiveness and stability of the method. Finally, the processing and analysis in this study involved the collection of nanopore sequencing data from the published literature, but the actual clinical samples were not extracted and analyzed. In the future, we plan to collect over 200 clinical samples, covering different pathogens. In general, this platform represents the latest application and exploration of nanopore sequencing in the field of rapid detection of pathogenic microorganisms and provides a new tool for microbial pathogen or AMR detection research.

## Acknowledgements

We thank Fujian Susuzhao Technology Co., Ltd. for providing technical support for the development of algorithm online prediction platform.

## Authors’ contributions

G.M. conceived the project, designed the experiment, analyzed data, and wrote the manuscript. Y.D. collected and analyzed data. All authors reviewed the manuscript.

## Availability of data and materials

All data generated or analyzed during this study are included in this published article and in the GitHub.

## Consent to Publish

Not applicable.

## Figure Legends

Supplementary Table 1. Detection results of sample S6 with detailed information of pathogen identification, including six parameters, namely number_of_matches, average, sum, reads_ratio, all_sum, and all_ratio.

Supplementary Table 2. Detection results of sample S6 with detailed information of antibiotic-resistance gene identification, including six parameters, namely number_of_matches, average, sum, reads_ratio, all_sum, and all_ratio.

Supplementary Table 3. Detection results of sample S13 with detailed information of pathogen identification, including six parameters, namely number_of_matches, average, sum, reads_ratio, all_sum, and all_ratio.

Supplementary Table 4. Detection results of sample S13 with detailed information of antibiotic-resistance gene identification, including six parameters, namely number_of_matches, average, sum, reads_ratio, all_sum, and all_ratio.

Supplementary Table 5. Detection results of sample E1 with detailed information of pathogen identification, including six parameters, namely number_of_matches, average, sum, reads_ratio, all_sum, and all_ratio.

Supplementary Table 6. Detection results of sample E2 with detailed information of pathogen identification, including six parameters, namely number_of_matches, average, sum, reads_ratio, all_sum, and all_ratio.

